# Induction of Human Pruriceptors from Pluripotent Stem Cells via Transcription Factors

**DOI:** 10.1101/2025.06.11.659208

**Authors:** Hisato Iriki, Ruiqi Hu, Xu Li, Erdene Baljinnyam, Carina Habich, Ichiro Imanishi, Loan Miller, Kavya Chegireddy, Laraib Iqbal Malik, Daniel Yassky, Aaron Ver Heul, Kathleen M. Smith, Eric R. Goedken, Peter Reinhardt, Brian S. Kim, Samuele G. Marro

**Affiliations:** Nash Family Department of Neuroscience, Friedman Brain Institute, Icahn School of Medicine at Mount Sinai, New York, NY, USA; Institute for Regenerative Medicine, Icahn School of Medicine at Mount Sinai, New York, NY, USA; Kimberly and Eric J. Waldman Department of Dermatology, Icahn School of Medicine at Mount Sinai, New York, NY, USA; Mark Lebwohl Center for Neuroinflammation and Sensation, Icahn School of Medicine at Mount Sinai, New York, NY, USA; Marc and Jennifer Lipschultz Precision Immunology Institute, Icahn School of Medicine at Mount Sinai, New York, NY, USA; AbbVie Bioresearch Center, Worcester, MA, USA; AbbVie Inc., North Chicago, IL, USA; AbbVie Deutschland GmbH & Co KG, Neuroscience Discovery, Knollstrasse, Ludwigshafen am Rhein, Germany; Allen Discovery Center for Neuroimmune Interactions, Icahn School of Medicine at Mount Sinai, New York, NY, USA; Division of Allergy and Immunology, John T. Milliken Department of Medicine, Washington University School of Medicine in St. Louis, 660 S. Euclid Avenue, MSC 8122-0021-03, St. Louis, MO, USA

**Keywords:** hPSCs, Pruriceptors, NGN1, ISL1, JAK1-inhibitor, iPruriceptors

## Abstract

Pruriception is crucial for defense against external stimuli but can lead to chronic pruritus, a debilitating condition affecting millions worldwide. Our understanding of the cellular and molecular mechanisms behind the sensation of itch has been hindered by the lack of functional human models. Here, we address this limitation by developing a protocol to generate induced pruriceptors (iPruriceptors) from human pluripotent stem cells (hPSCs). We compared two differentiation approaches: a direct method via forced expression of transcription factors (TFs) in hPSCs, and a 2-step process through expression of TFs in hPSC-derived neural crest cells (NCCs). The 2-step protocol proved superior in inducing a transcriptional program that closely resembles that of human pruriceptors. Our optimized protocol employs forced expression of NGN1 and ISL1 to drive differentiation from NCCs into pruriceptors, enhancing the expression of known pruritogen receptors such as IL31RA, which pairs with OSMR, and HRH1. The induction of this transcriptional program leads to functional maturation of iPruriceptors. Accordingly, iPruriceptors exhibit robust responses to itch stimuli and *in vivo*-like itch pharmacology such as treatment with ABT-317, a JAK1 inhibitor tool compound, similar to those targeting intensive pruritus in atopic dermatitis (AD). Importantly, iPruriceptors can be generated without viral vectors or safe-harbor gene editing, using a PiggyBac-based transfection method that simplifies scalability. Our protocol offers a robust platform for investigating itch biology, modeling chronic pruritus, and enabling high-throughput screening for therapeutic target discovery.

**Highlights:**

- NGN1 and ISL1 forced expression in NCCs induces rapid differentiation to iPruriceptors
- iPruriceptors share transcriptional profile of primary human pruriceptors
- iPruriceptors have electrophysiological responses to known pruritogens
- iPruriceptors have JAK1-dependent IL-31/-13 responses blocked by ABT-317

## Introduction

Pruriception, the perception of itch, helps the body defend against parasites and environmental irritants by provoking the desire to scratch. However, excessive and prolonged itch can lead to chronic pruritus, defined as itch lasting more than six weeks. Chronic pruritus affects over seven million people annually in the United States alone and can arise from multiple underlying causes, including immunological factors, systemic diseases, neuropathic origins, infections, drug-induced reactions, and/or idiopathically as in chronic pruritus of unknown origin (CPUO), among others (Butler *et al*., 2024; Kim *et al*., 2019).

The molecular mechanisms of itch begin with a stimulus in the skin that triggers local cells to produce cytokines and mediators or through direct activity on itch-sensory neurons called pruriceptors. Stimuli that elicit itch by acting directly on key molecular receptors on pruriceptors are referred to as pruritogens. Once activated, these receptors initiate a complex signaling cascade (Chen *et al*., 2020), leading to neuronal depolarization, action potential generation, and neurotransmitter release. This signal is then transmitted along the spinothalamic tract to the brainstem, where it is processed and relayed to various brain regions, ultimately producing the urge to scratch (Chen *et al*., 2020; Wang *et al*., 2021).

A well-known pruritogen is histamine, which mediates acute itch by activating the HRH1 receptor (histaminergic pathway) upon release from mast cells (Yosipovitch *et al*., 2018). Other mediators include IL-4, IL-13, and IL-31, type 2 cytokines produced by lymphocytes and mast cells, which activate their receptors for stimulating non-histaminergic itch pathways. Other canonical markers of pruriceptors include PAR-2, CysLTR2, somatostatin (SST), and B-type natriuretic peptide (BNP). Among these, BNP serves as a key neurotransmitter that mediates itch transmission to the spinal cord by activating NPR-A receptors. Notably, mice lacking BNP or NPR-A exhibit reduced responses to multiple itch stimuli, highlighting its crucial role in itch signaling (Meng *et al*., 2021; Mishra and Hoon, 2013). Common itch-associated cytokine receptors signal through the Janus kinase/signal transducer and activator of transcription (JAK/STAT) (Oetjen *et al*., 2017). Patients with recalcitrant chronic itch that have failed other immunosuppressive therapies have shown marked improvement when treated with Janus kinase 1 (JAK1)-selective inhibitors (Huang *et al*., 2024; Jeon *et al*., 2022; Kim *et al*., 2021; Kim *et al*., 2020). Indeed, JAK1 is highly expressed in itch neurons in human dorsal root ganglia (DRG) sensory neurons (Nguyen *et al*., 2021). Beyond canonical markers, recent single cell multiomic studies of mouse and human DRG have expanded and reclassified neuronal subtypes across species. Interestingly, pruriceptors are among the most divergent groups between humans and mice. One study identified new classes of human pruriceptors named H10 and H11, which share some features with new classes of mouse pruriceptors named NP1 and NP3 (Nguyen *et al*., 2021; Usoskin *et al*., 2015). However, a unique feature of human pruriceptors is that they exhibit many more non-pruriceptive receptors, suggesting their capacity for functional polymodality (Jung *et al*., 2023). Thus, the ability to functionally evaluate human sensory neurons would greatly advance clinical development of novel anti-pruritic therapies.

Developmentally, pruriceptors are a subclass of nociceptors, specifically belonging to the non-peptidergic (NP) class, in contrast to the peptidergic (PEP) class. Like other nociceptors, pruriceptors originate from neural crest cells (NCCs) or, in the case of trigeminal nociceptors, from the head placode. Nociceptors are characterized by the expression of TrkA (tropomyosin receptor kinase A), the high-affinity receptor for nerve growth factor (NGF), and are generated during the last wave of sensory neuron development (Ma *et al*., 1999; Sommer *et al*., 1996; Zirlinger *et al*., 2002). Proprioceptors are the earliest to develop and express TrkC, the receptor for neurotrophin-3 (NT-3). Mechanoreceptors follow, expressing TrkB, the receptor for brain-derived neurotrophic factor (BDNF). The first two waves of sensory neuron development (proprioceptors and mechanoreceptors) are driven by Neurogenin-2 (NGN2), while the third wave (nociceptors including pruriceptors) is controlled by Neurogenin-1 (NGN1) (Marmigere and Ernfors, 2007).

Numerous protocols have successfully recapitulated this complex developmental cascade *in vitro*, enabling differentiation of human pluripotent stem cells (hPSCs) into sensory neurons using either extrinsic factors, forced expression of lineage-specifying transcription factors (TFs), or a combination of both. For instance, hPSCs have been directed toward NCCs and subsequently induced to adopt a sensory neuron fate through the forced expression of TFs such as NGN1 or NGN2. The resulting sensory neurons exhibit characteristics of general nociceptors (Holzer *et al*., 2022; Lallemend and Ernfors, 2012; Marmigere and Ernfors, 2007; Nickolls *et al*., 2020; Plumbly *et al*., 2022; Schrenk-Siemens *et al*., 2022). However, it has been reported that there is no difference in nociceptor induction efficiency between NGN1 and NGN2, at least in forced expression in fibroblasts (Blanchard *et al*., 2015), and the significance of NGN1 over NGN2 in NCC and the ideal use of these TFs has not been clarified completely. In addition, it is known that other TFs are also expressed in discrete stages during nociceptors differentiation. How they work in conjunction with NGN1 and NGN2 in the differentiation of hPSCs is largely unknown.

To date, no protocol has been established for specifically generating pruriceptors from hPSCs through the expression of lineage-specifying TFs. Here, we describe a novel protocol that generates *bona fide* human pruriceptors from hPSCs through the forced expression of lineage-specifying TFs. The resulting cells (iPruriceptors) express markers and receptors for pruritogens identified in primary DRG single-cell transcriptomic studies. Using a stepwise protocol that mimics *in vivo* sensory neuron differentiation, we confirmed their responsiveness to itch stimuli and the ability of a JAK1-selective inhibitor to disrupt their activity in line with the known anti-pruritic effects of clinical JAK1 inhibition. This approach provides a reliable, reproducible and scalable method for generating human pruriceptors, offering new insights into pruriception and potential therapeutic applications for chronic itch disorders.

## Results

### Enhanced pruriceptor fate induction by NGN1 overexpression in NCCs

Protocols for the differentiation of hPSCs to neural lineage based on extrinsic factors faithfully reproduce the stepwise processes of neurogenesis but are known to be inefficient and lengthy (Chambers *et al*., 2009; Mica *et al*., 2013). We therefore turned to the widely used ectopic expression of the master regulator of neurogenesis, Neurogenin-2 (NGN2), to bypass the progenitor state and rapidly differentiate hPSCs to induced neurons (iN) with minimal line-to-line variability (Zhang *et al*., 2013). To optimize the specification of iN cells toward pruriceptors, we focused on a panel of markers typically expressed by primary human pruriceptors as defined by the new classes H10 and H11(Nguyen *et al*., 2021). In parallel to NGN2, we also tested Neurogenin-1 (NGN1) because of its role in the neurogenesis of pruriceptors (Ma *et al*., 1999; Sommer *et al*., 1996; Zirlinger *et al*., 2002). NGN1 has been used to generate motor neurons from hPSCs (Goparaju *et al*., 2017), mixed population of sensory neurons from fibroblasts (Blanchard *et al*., 2015) and hPSCs (Holzer *et al*., 2022). We therefore asked whether NGN2 and NGN1 have different effects when overexpressed in hPSC (1-step protocol) or in hPSC-derived NCCs (2-step protocol). For the 1-step protocol, we used lentiviral transduction for constitutive expression of rtTA (Urlinger *et al*., 2000) and tetracycline-inducible expression driven by a Tetracycline (Tet) inducible promoter to express either NGN1 or NGN2 in hPSCs, followed by doxycycline induction on day 1 and positive selection with puromycin on day 2. For the 2-step protocol, we first differentiated hPSCs into NCCs using the LSB3i protocol previously reported (Chambers *et al*., 2012). At day 11, we transduced the NCCs with the same lentiviral constructs to express either NGN1 or NGN2, followed by doxycycline induction on day 12 and positive selection with puromycin (Puro) on day 13 (Figure 1A). At day 21 both protocols produced bipolar morphology typical of iN cells (Figure 1B). We then analyzed the transcriptomic profile of the resulting iN cells using RNA sequencing (RNA-seq) and confirmed that pluripotency markers such as POU5F1, and NANOG were efficiently downregulated in both protocols (Figure 1C, and S1B).

**Figure 1.**
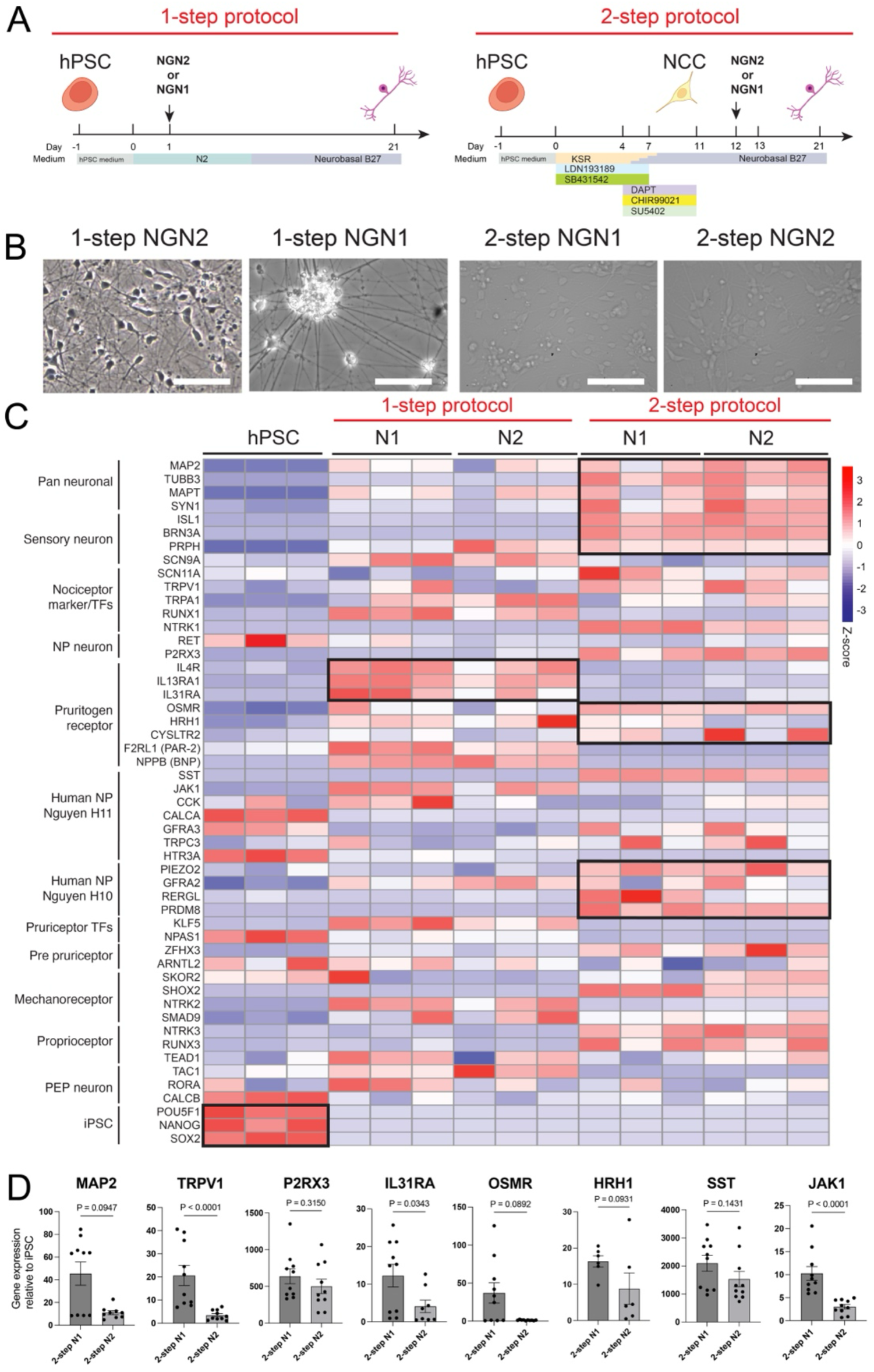
NGN1 expression in NCCs shows a tendency to enhance pruriceptor marker induction. (A) Schematic of NGN1 and NGN2 expression in hPSCs (1-step protocol) and NCCs induced by LSB3i (2-step protocol) using a lentiviral system. (B) Representative images showing neuronal morphology in 1-step NGN1, 1-step NGN2, 2-step NGN1, 2-step NGN2 on Day 21. Scale bars indicate 100μm. (C) Heatmap analysis of samples obtained on protocol day 21 using either the 1-step or 2-step protocol with NGN1 or NGN2 expression (1-stepN1, 1-stepN2, 2-stepN1, and 2-stepN2), along with hPSCs, showing the expression of the indicated markers. Transcript expression values (TPM) were log2-transformed, and Z-scores were calculated, n=3. (D) Quantitative PCR analysis of indicated markers in 2-step N1 and 2-step N2 protocol on day 21. Relative expression levels were normalized to control hPSC levels. Data are presented as mean ± SEM, with individual data points representing independent biological replicates. Statistical comparisons were conducted using an unpaired t-test. The data are pooled from two independent experiments.

Interestingly, the 2-step protocol more effectively induced pan-neuronal markers such as MAP2 and MAPT, indicating a more robust promotion of neuronal differentiation. Notably, the choice of protocol (1-step vs. 2-step) had a greater impact on differentiation outcomes than the identity of the overexpressed TF as shown by the principal component analysis (Figure S1A). Within the 1-step protocol, however, NGN1 induced higher levels of nociceptor-associated genes, including RUNX1 and TRPV1, relative to the NGN2 protocol. Similarly, NGN1 outperformed NGN2 in promoting the expression of pruritogen receptors and associated signaling components such as IL13RA1, IL31RA, OSMR, HRH1, and JAK1 (Figure 1C and S1B). For H10 markers, the 2-step protocol—particularly in combination with NGN1—also showed a distinct advantage. Similarly, the expression of HRH1, one of the most representative pruritogen receptors, was higher in 2-step NGN11 than in 2-step NGN2.

In a separate differentiation batch, we performed quantitative real-time PCR (qPCR) to directly compare the efficiency of NGN1 and NGN2 in the two-step protocol for inducing the expression of key genes, including IL31RA, OSMR, and JAK1. Overall, NGN1 demonstrated superior performance (Figure 1D).

Together, these results suggest that NGN1 has a distinct capacity to initiate specific transcriptional programs compared to NGN2, and that the execution of these programs is influenced by both the cellular context in which NGN1 is overexpressed and the extrinsic signals present during induction. Given the superior efficiency of the 2-step protocol in promoting neuronal gene expression and suppressing the NCC program, along with the enhanced induction of pruriceptor markers, we selected the 2-step NGN1 protocol for subsequent refinement.

### ISL1 contributes to DRG cell fate specification when co-expressed with NGN1

To further refine a protocol for the generation of iN that faithfully represent human pruriceptors, we added additional TFs that are involved in the second wave neurogenesis, namely ISL1, BRN3A, and RUNX1 to the NGN1-2-step overexpression protocol (Lallemend and Ernfors, 2012; Marmigere and Ernfors, 2007) (Figure 2A,B). These refined protocols—named 2-step N1-I, 2-step N1-B, and 2-step N1-R—produced iN cells with the same apparent efficiency of the 2-step N1 protocol with no effect on morphology. We then analyzed the effect of the added TFs on the global transcription profile of the iN. Not surprisingly, we observed some level of cross-induction between the TFs overexpressed in the different protocols, for instance, in the N1 protocol NGN1 induced the expression of NGN2 (to the same levels found in the N2 protocol), but in the N2 protocol NGN2 only weakly induced NGN1 (Figure S2A). Also, ISL1 expression in the context of N1-I protocol synergized with NGN1 to further induce NGN2 expression, and the expression of BRN3A. On the contrary, no cross-induction was observed in the N1-B or N1-R protocols (Figure 2C). On a global scale, the transcriptional profile of the iN was majorly altered by the addition of BRN3A or RUNX1, as indicated by the unsupervised hierarchical clustering of variance-stabilized counts (Pearson-correlation distance, Ward’s linkage) of the N1-B and N1-R in a separate group from the N1, N2 and N1-I (Figure 2D).

**Figure 2.**
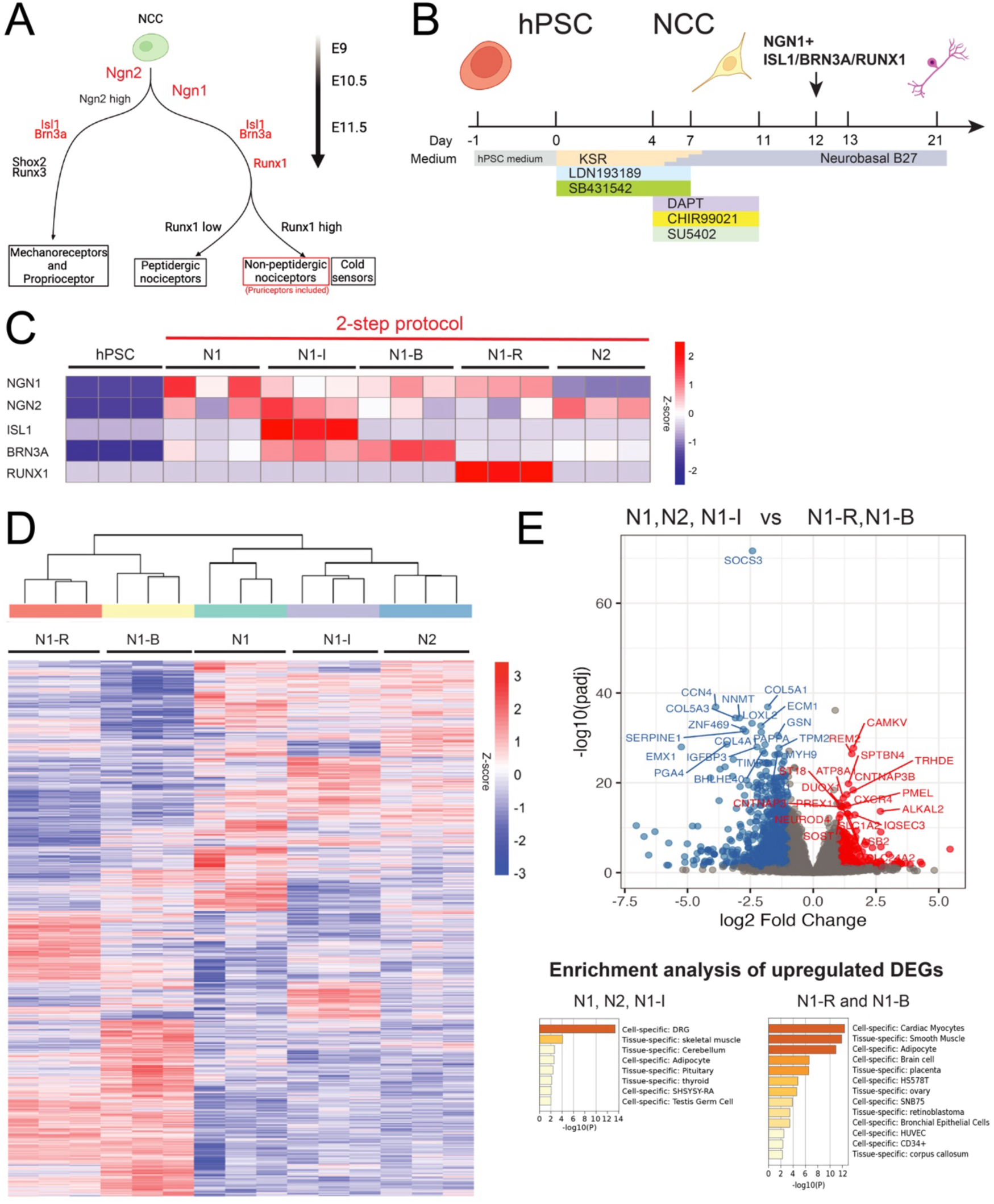
ISL1 contributes to differentiation into DRG neurons when co-expressed with NGN1. (A) Schematic of peripheral nerve developmental cascade in mice highlighting the role of NGN1, ISL1, BRN3A, and RUNX1 in NP-nociceptor differentiation. (B) Schematic of co-expression of NGN1 with ISL1, BRN3A, and RUNX1 expression in NCCs induced by LSB3i using a lentiviral system. (C) Heatmap analysis of samples obtained on protocol day 21 using 2-step N1, N1-I, N1-B, N1-R, and N2, along with hPSCs, showing the expression of the introduced TFs; NGN1, NGN2, ISL1, BRN3A, and RUNX1. Transcript expression values (TPM) were log10-transformed, and Z-scores were calculated. (D) Heatmap analysis and hierarchical clustering using the expression of DEGs on the samples from 2-step N1, N1-I, N1-B, N1-R, and N2. (E) Volcano plot displaying differentially expressed genes (DEGs) identified by comparing the N1 / N2 / N1-I group with the N1-R / N1-B group (all samples derived from the 2-step protocol). The x-axis shows log2 fold change and the y-axis shows –log10 (adjusted P value). Genes with an adjusted P value < 0.05, calculated with DESeq2, were considered significantly up- or down-regulated. The bar charts below summarise Metascape enrichment (–log_10_ P) of the top 300 up-regulated genes: left, tissues/cell types over-represented in N1 / N2 / N1-I; right, those over-represented in N1-R / N1-B.

We then performed differentially expressed gene (DEG) analysis of these two groups and observed that only the second group (N1, N2 and N1-I) expressed genes associated to general sensory neuron identity such as ALKAL2 and REM2 (Defaye *et al*., 2022; Liput *et al*., 2016) (Figure 2E and S2B, Table S1). On the other hand, the first group (N1-B and N1- R) had robust expression of non-PNS markers (Figure 2E, Table S1). Accordingly, N1-I showed high expression of genes related to PNS and DRG neurons when it was compared with N1-B or N1-R (Figure S2B and Table S2). Together, these results suggest that ISL1— but not BRN3A and RUNX1— when co-expressed with NGN1, contributes to the activation of a transcriptional program resembling that of DRG neurons.

In addition to enhancing a DRG-like transcriptional program, the combination of NGN1 and ISL1 also promoted the expression of pruritogen receptors. Compared to NGN1 alone, N1-I iN cells exhibited stronger expression of peripheral nervous system (PNS) markers (PRPH), nociceptor markers (RET, P2RX3), pruritogen receptors and associated signaling proteins (e.g., JAK1), as well as markers of the H10 and H11 neuronal subclasses (Figure 3A). Based on these findings, we concluded that co-expression of NGN1 and ISL1 in NCC drives pruriceptor specification. We therefore named these cells ‘iPruriceptors’ due to their rapid neuronal differentiation (Figure 3B) by day 12, and their robust expression of pruriceptor markers—including RET, IL31RA, TRKA, TRPV1, and OSMR—by day 28, as confirmed by immunohistochemistry (Figure 3C).

**Figure 3.**
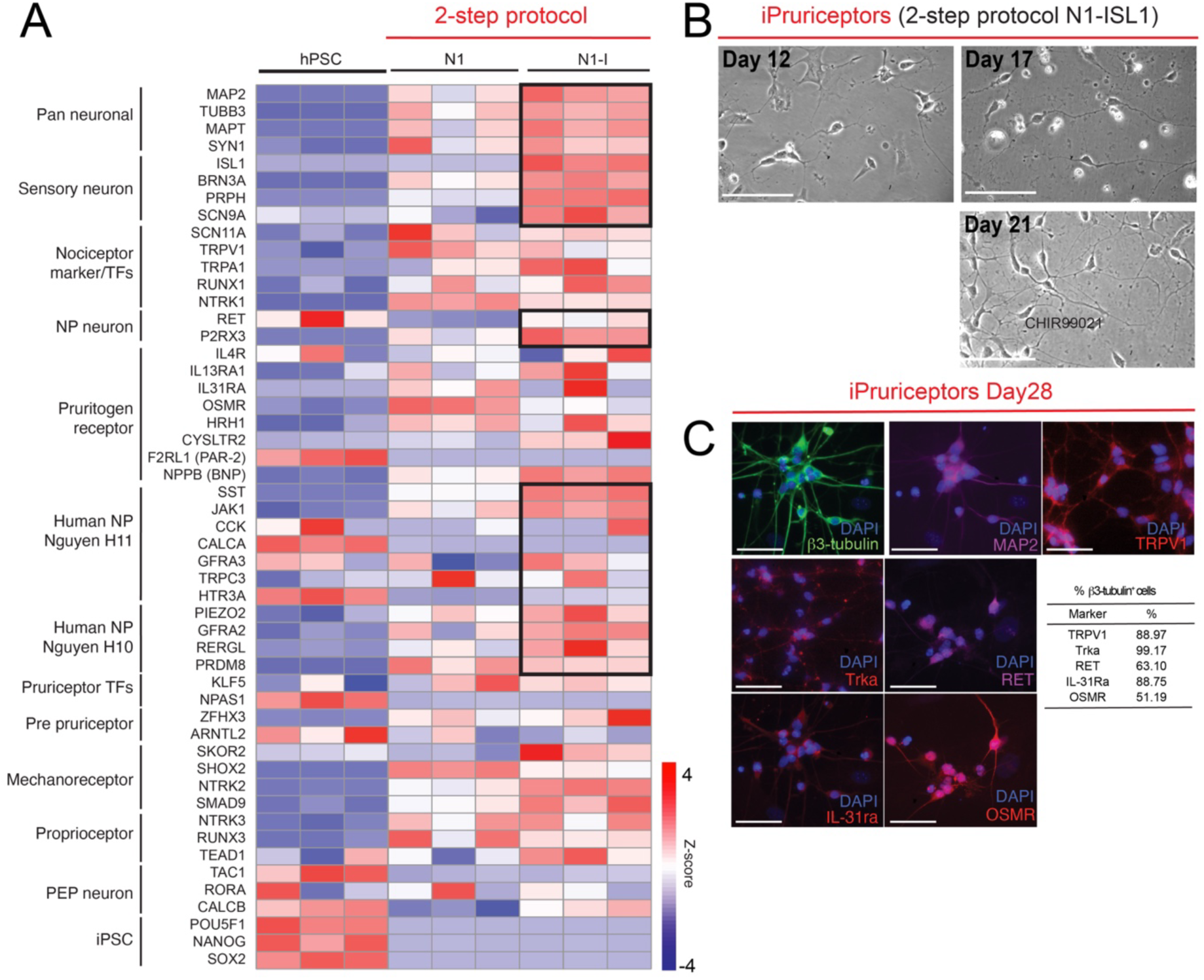
Co-expression of NGN1 and ISL1 induce pruriceptor marker expression. (A) Heatmap analysis on the samples from 2-step N1 and N1-I, showing the expression of indicated markers. Transcript expression values (TPM) were log10-transformed, and Z-scores were calculated. (B) Representative images showing neuronal morphology at different stages of differentiation under the 2-step N1-I, namely iPruriceptor, protocol on day12, 17, 21. Scale bars indicate 100μm. (C) Immunofluorescence staining of indicated markers on the induced neurons under iPruriceptor protocol on day 28. Scale bars indicate 50μm. The table shows the proportion of induced neurons under iPruriceptor protocol expressing the indicated markers on day 28. The percentages of TRPV1-, TrkA-, RET-, IL31ra-, or OSMR-positive cells among the β3 tubulin–positive population, which were stained simultaneously with each described marker, were shown.

### iPruriceptors selectively respond to pruritogenic stimuli and JAK1 inhibition

Next, we sought to determine whether iPruriceptors are electrophysiologically active and capable of responding to specific pruritogenic stimuli, such as histamine and IL-31, as well as to general, non-pruritogenic agents like capsaicin which signals via TRPV1. To promote their functional maturation, we co-cultured iN cells with primary mouse glial cells starting on day 18, following our previously established iN protocols (Marro *et al*., 2019; Pang *et al*., 2011; Zhang *et al*., 2013). The culture medium of iN cells is typically supplemented with BDNF, the primary ligand of TrkB, a classical marker of mechanoreceptors (Lu *et al*., 2024). We therefore tested the effect of supplementing the maturation medium with NGF, the ligand for TrkA, a key marker of nociceptors, and asked whether NGF—administered during the maturation phase rather than during initial induction—could enhance the electrophysiological properties of iPruriceptors (Figure S4A).

Importantly, because this supplementation occurred after the transcriptional profile of the cells was presumably already established, we predicted a negligible impact on overall cell fate. Nevertheless, to exclude this possibility, we assessed the expression of key pruriceptor marker genes under both conditions and found that all expected markers remained robustly expressed—with some even upregulated in NGF-supplemented cultures—suggesting a robust differentiation into pruriceptors (Figure S4B). We next compared the two maturation conditions—BDNF vs. NGF—at the functional level using multielectrode array (MEA) recordings (Figure 4A and B). iPruriceptors matured in NGF-containing media exhibited a lower basal activity compared to those matured in BDNF, as measured by mean firing rate (MFR) (Figure 4C, S4C). BDNF cultures also displayed more frequent network bursts (Figure S4D and E), indicative of synchronized firing often observed in glutamatergic CNS neurons.

**Figure 4.**
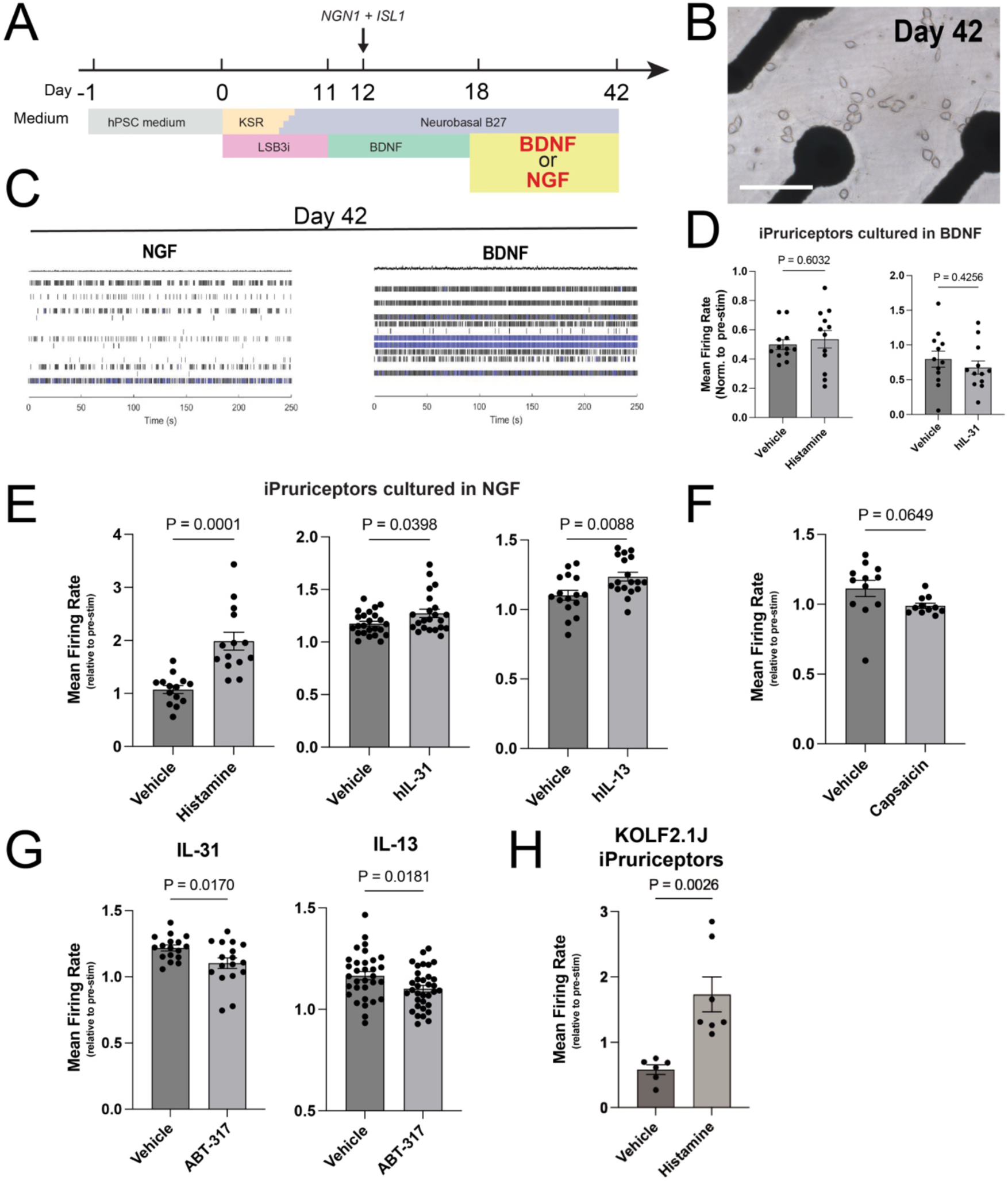
iPruriceptors functionally respond to pruritogenic stimuli and JAK1 inhibition. (A) Schematic showing co-expression of NGN1 and ISL1 in NCCs induced by LSB3i via lentiviral transduction, followed by culture with BDNF or NGF from day 18 to day 42, the time point of MEA analysis. (B) Representative images showing neuronal morphology under iPruriceptor protocol on day 42 on the MEA plate. Scale bars indicate 100 μm. (C) MEA raster plots showing neuronal spiking activity of neuronal activity on day 42 under different neurotrophic conditions (BDNF/GDNF vs NGF) after day 18. Each vertical line represents an action potential recorded from a single electrode. (D) MEA analysis showing neuronal responses to histamine and IL-31 (pooled from two independent experiments, n = 6 each) in induced neurons after day 42 from the iPruriceptor protocol maintained with BDNF/GDNF after day 18. (E) MEA analysis showing neuronal responses to histamine (pooled from two independent experiments, n = 6 each), IL-31 (pooled from four independent experiments, n = 6 each), and IL-13 (pooled from two independent experiments, n = 6–12 each) after day 42 in induced neurons from the iPruriceptor protocol maintained with NGF after day 18. (F) MEA analysis showing neuronal responses to capsaicin (pooled from two independent experiments, n = 6 each) on day 42 in induced neurons from the iPruriceptor protocol maintained with NGF after day 18. (G) MEA analysis showing firing rates before and after IL-31 (pooled from three independent experiments, n = 6–8 each) and IL-13 (pooled from three independent experiments, n = 10–12 each) after day 42 in the presence or absence of a JAK1 inhibitor, ABT-317. (H) MEA analysis showing neuronal responses to histamine in induced neurons, using KOLF2.1 J cell line, on day 42 from the iPruriceptor protocol maintained with NGF after day 18. Representative data from two independent experiments (n = 6 or 7). As for MEA analysis, Values represent mean firing rates normalized to pre-stimulation baseline—per well in (D) - (G), and to the average baseline across wells per condition in (H). Data are presented as mean ± SEM. Each dot represents one well. Statistical comparisons between the two groups were performed using an unpaired t-test with Welch’s correction.

To evaluate stimulus-induced activity, we exposed iPruriceptors to histamine and IL-31. iN cells matured in BDNF-containing media showed no change in mean firing rate (MFR) following stimulation (Figure 4D). In contrast, iPruriceptors cultured with NGF exhibited a significant increase in MFR in response to histamine, IL-31, and IL-13 (Figure 4E).

Treatment with the non-pruritogenic agent capsaicin did not elicit any electrophysiological response in either condition, confirming the specificity of the response (Figure 4F).

Finally, we tested whether the IL-31 and IL-13 responses were mediated by JAK1 signaling using the JAK1 inhibitor tool compound ABT-317 (Ge *et al*., 2020; Todorovic *et al*., 2022; Twomey *et al*., 2024). Inhibition of neuronal JAK1 pathways has also been shown to ameliorate pruritus (Oetjen *et al*., 2017). Remarkably, this treatment fully suppressed the cytokine driven MEA response, indicating that iPruriceptors respond specifically via the JAK1-dependent pathway (Figure 4G), consistent with the expected anti-pruritogenic properties. To extend the validation of these results and test the robustness of the protocol across a broader range of iPSC lines, we generated iPruriceptors from one additional hPSC line, the KOLF2.1J (Pantazis *et al*., 2022). KOLF2.1J produced iPruriceptors with similar efficiency of our reference line (WTC11), showing the pruriceptor marker expression on day 21 (Figure S4F). Additionally, the resulting iPruriceptors were functionally active, as shown by their response to histamine (Figure 4H). These functional data support the utility of iPruriceptors as a relevant *in vitro* platform for translational studies of pruritus.

Finally, to enhance the scalability of the iPruriceptors protocol across multiple hPSC lines, we sought alternatives to lentiviral delivery of NGN1 and ISL1, a common bottleneck in iN cell production. As an alternative, we employed the PiggyBac (PB) transposon system, previously used for iN cell generation (Gu *et al*., 2022). We constructed a polycistronic PB vector containing: (1) the reverse transactivator rtTA under the control of CAG promoter; (2) a puromycin resistance gene and a nuclear-localized BFP under the control of EF1α promoter; (3) a ubiquitous chromatin opening element (UCOE) to attenuate transgene silencing in hPSCs and their differentiated cells (Durens *et al*., 2024; Muller-Kuller *et al*., 2015); and (4) a bidirectional doxycycline-inducible TRE3G BI promoter to drive co-expression of NGN1 and ISL1 (Figure 5A). Following PB transfection, hPSCs expressed nuclear BFP and differentiated into iN cells at a rate comparable to those generated using lentiviral vectors (Figure 5B). Immunohistochemistry confirmed robust expression of β3-tubulin, MAP2, RET, IL-31RA, and OSMR at day 28 (Figure 5C). Finally, MEA recordings revealed functional responses to histamine and hIL-31 (Figure 5D). Together, these results demonstrate that functional iPruriceptors can be efficiently generated from hPSCs using a transfection-based method and without viral vectors.

**Figure 5.**
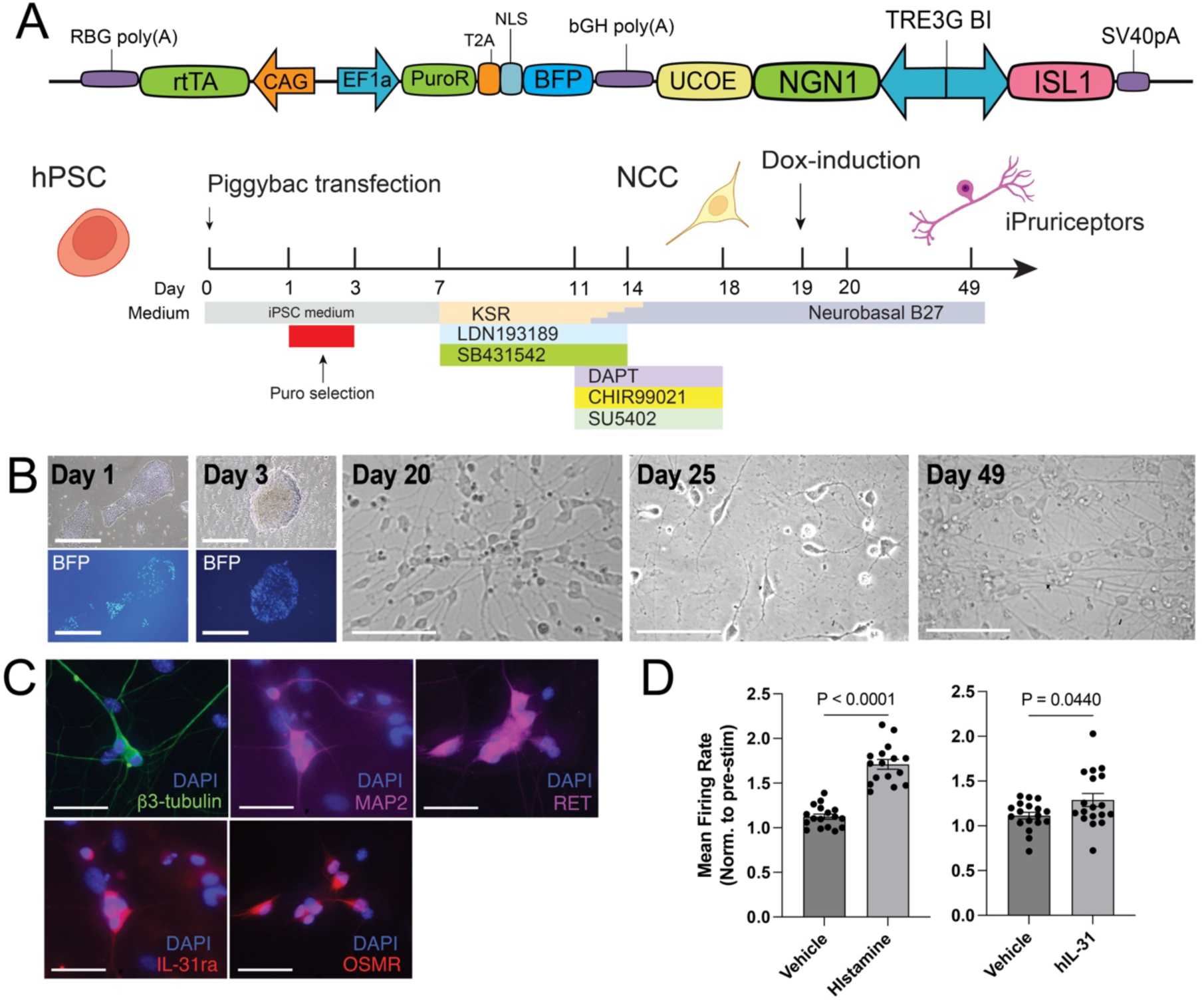
iPruriceptor with PiggyBac system reproduce robust functional pruriceptor induction. (A) Schematic of co-expression of NGN1 with ISL1 in NCCs induced by LSB3i using PiggyBac system, PB-iPruriceptor protocol. (B) Representative images showing neuronal morphology and BFP expression at different stages of differentiation under the PB-iPruriceptor protocol on day 1, 3, 20, 25, 49. Scale bars indicate 100 μm. (C) Immunofluorescence staining of indicated markers on the induced neurons under PB-iPruriceptor protocol on day 35. DAPI staining are shown in blue and the slightly larger nuclei that are stained granularly are co-plated CD-1 IGS mouse glial cells. Scale bars indicate 50 μm. (D) MEA analysis showing histamine-, and IL-31-induced neuronal responses on induced neurons from PB-iPruriceptor protocol on day 49. The values represent the mean firing rate, normalized to the baseline firing rate before stimulation for each well. Data are presented as mean ± SEM. Each dot represents one well. Data is pooled from three independent experiments, n = 4–12 each). Statistical comparisons between the two groups were performed using an unpaired t-test with Welch’s correction.

## Discussion

To our knowledge, this is the first study reporting the induction of pruriceptors from hPSCs through the forced expression of lineage-specifying TFs. A previous study described the generation of itch sensory neuron-like cells (ISNLCs) from hPSCs using a method based solely on extrinsic factors (Guo *et al*., 2022). While ISNLCs express pruriceptor markers, they exhibit variability in their responses to histamine and PAR-2, lack expression of the IL- 31 receptor (IL31RA), and the overall neuronal induction efficiency was not clearly quantified. Another study using extrinsic factor-based differentiation reported up to 90% efficiency in generating sensory neurons expressing IL31RA; however, functional characterization was limited to calcium influx measurements (Umehara *et al*., 2020).

Our protocol integrates the precision of extrinsic factors patterning with the speed and efficiency of TF–mediated reprogramming to generate iPruriceptors that are both molecularly and functionally similar to primary human pruriceptors. By inducing TFs in NCCs, rather than directly in hPSCs, our approach enhances culture synchronization and promotes neuronal maturation, achieving approximately 77% efficiency (defined as the proportion of βIII-tubulin+, MAP2+, and IL31RA+ cells produced by the end of protocol). The forced expression of TFs has been employed by many groups as tool to differentiate pluripotent or somatic cells into iN (Pang *et al*., 2011), for instance the basic helix–loop– helix (bHLH) NGN2 has been widely used as versatile neurogenic TF—for a comprehensive review see Hulme et al. (Hulme *et al*., 2022)— to derive iN with features of CNS and PNS neurons including sensory neurons (Plumbly *et al*., 2022).

Our approach to generate iN with pruriceptor features (iPruriceptors) is based on the observation that mouse DRG sensory neuron development is initiated by a neurogenic phase induced by TFs of the neurogenin class (Fode *et al*., 1998; Ma *et al*., 1999). Both NGN1 and NGN2 and play a critical neurogenic role, as double knockout mice fail to develop DRGs.

However, these two factors have distinct roles: NGN2 is specifically required for the generation of mechanoreceptive and proprioceptive TrkC+ and TrkB+ neurons, whereas the formation of nociceptive TrkA+ neurons relies exclusively on NGN1 (Ma *et al*., 1999).

Following the neurogenic phase, sensory neurons initiate expression of the homeodomain TFs, Brn3a and Isl1 (Eng *et al*., 2001; Fedtsova and Turner, 1995; Sun *et al*., 2008).

Although Brn3a (a POU domain factor) and Islet1 (a LIM-homeodomain factor) belong to distinct TF families, they have overlapping downstream targets. Their co-expression in all sensory neurons has led to the proposal that they constitute a combinatorial code for sensory lineage specification (Anderson, 1999; Dykes *et al*., 2011).

Finally, Brn3a and Islet1 synergistically interact with TFs of the RUNX family to establish the gene regulatory program required for the formation of TrkC+ neurons (Orlovsky *et al*., 2024).

In line with these developmental roles, we found that ISL1, when co-expressed with NGN1 in NCCs, robustly induces a pruriceptor-specific transcriptional program, partly through the upregulation of BRN3A. However, the reverse is not true: BRN3A does not induce ISL1 expression, nor does it efficiently drive sensory neuron fate on its own. This finding is surprising, particularly because BRN3A has previously been used in combination with NGN2 to generate mixed populations of DRG-like sensory neurons from fibroblasts, hPSCs, and NCCs (Blanchard *et al*., 2015; Nickolls *et al*., 2020). A likely explanation lies in the context-dependent functions of BRN3A, which may act in concert with neuron subtype– specific cofactors. In fact, BRN3A is known to function as a terminal selector of neuronal identity in the CNS (Serrano-Saiz *et al*., 2018), and a hierarchical relationship between ISL1 and BRN3A has also been observed during inner ear neuron development, where ISL1 expression precedes and possibly regulates BRN3A (Deng *et al*., 2014; Xu *et al*., 2024).

Similarly, the observation that RUNX1 is inefficient at inducing iPruriceptors is unexpected but can be explained by its position downstream of ISL1 and BRN3A in the transcriptional hierarchy. RUNX1 is known to refine nociceptor subtype identity rather than initiate sensory lineage specification. When co-expressed with an early neurogenic factor like NGN1, RUNX1 may fail to activate an appropriate pruriceptor gene expression program.

This is consistent with its known developmental dynamics: RUNX1 is initially activated by ISL1 and BRN3A but later represses early sensory markers such as TrkA to promote subtype diversification (Chen *et al*., 2006). RUNX1’s dual role—acting first as a transcriptional activator and later as a repressor—adds complexity that may limit its utility in simplified *in vitro* differentiation models such as ours.

One key finding of this study is that pruriceptor differentiation is critically dependent on the NGF signaling environment. While most NGN2-iN models supplement their media with Brain-derived neurotrophic factor (BDNF) and Neurotrophin-3 (NT3) (Bullmann *et al*., 2024; Zhang *et al*., 2013), we observed that iPruriceptors only became functionally responsive to pruritogens when cultured in NGF-containing media, rather than BDNF. This aligns with *in vivo* studies demonstrating that NGF promotes nociceptive lineage maturation (Lu *et al*., 2024), while BDNF primarily supports mechanoreceptors and proprioceptors.

The requirement for NGF supplementation highlights the complex interplay between extrinsic neurotrophic signals and intrinsic transcriptional programs in sensory neuron differentiation.

A major strength of our study is the functional validation of iPruriceptors in a pharmacologically relevant context. Using MEA recordings, we demonstrated that iPruriceptors exhibit pruritogen-induced activity, which can be pharmacologically suppressed by the JAK1-selective inhibitor ABT-317. This finding has direct clinical relevance, as JAK1-selective inhibition is now a proven therapeutic strategy for chronic pruritus in multiple conditions (Kwatra *et al*., 2024; Simpson *et al*., 2024). The ability of our model to recapitulate drug responses observed in patients positions it as a valuable platform for drug discovery and mechanistic studies of itch-related disorders.

Furthermore, the PiggyBac transposon-based expression system offers an efficient and scalable method for generating iPruriceptors, bypassing the limitations associated with viral transduction. This innovation enhances the translational potential of our protocol, facilitating high-throughput screening of pruritic and anti-pruritic compounds.

## Limitations of study

While the current results show that our protocol reliably generates iPruriceptors, its validation remains limited to two hPSC lines, as such, its robustness across a broader range of hPSC lines has yet to be established. Another limitation is that the expression of pruriceptor markers has been investigated on a bulk population, a single-cell transcriptomic approach would provide finer resolution into cellular heterogeneity and subtype-specific differentiation efficiency. Also, while ISL1 co-expression improved pruriceptor specification, additional TFs, such as ETV1, ELF1, FOXO3, FOXN3 and KLF5 may further refine differentiation. Future studies should explore whether sequential expression of NGN1/ISL1 followed by these TF could enhance pruriceptor purity and functional maturation. Another key consideration is the absence of a fully reconstructed itch circuit *in vitro.* Although our model enables the study of peripheral neuron responses, future work should integrate co-culture systems with keratinocytes, immune cells, and spinal cord neurons to better mimic *in vivo* pruriceptive pathways. More studies will be necessary to further optimize and adapt this protocol for true high-throughput applications and to develop a robust drug screening platform suitable for preclinical interrogation of compounds targeting pruriceptor activity. A key goal will be the generation of iPruriceptors from a diverse panel of hPSC lines derived from both healthy individuals and patients with atopic dermatitis (AD) and chronic pruritus of unknown origin (CPUO). This will enable the identification of functional, disease-specific phenotypes and allow for personalized therapeutic targeting.

## Experimental Procedures

### hPSCs maintenance

The WTC11 cell line (Kreitzer *et al*., 2013) (hPSCreg UCSFi001-A) was obtained from the J. David Gladstone Institutes San Francisco, and the KOLF2.1J cell line (Pantazis *et al*., 2022) (hPSCreg WTSIi018-B-12) was obtained from the Jackson Laboratory (strain 765000). Both lines were cultured in 6-well plates coated with 1 mL of 60 µg/mL Geltrex (Thermo Fisher, A1413302) in DMEM/F12 (Thermo Fisher, 11320033) and maintained in hPSC medium (StemFlex, Thermo Fisher, A3349401). The medium was replaced every two days. When cultures reached ∼80% confluency, cells were gently detached by incubation with 0.5 mM EDTA in DPBS for 8 min at room temperature. Differentiation was assessed morphologically under an optical microscope before passaging. Detached cells were resuspended in hPSC medium supplemented with 2 µM Thiazovivin (Millipore Sigma, SML1045) and seeded onto new plates at split ratios between 1:10 and 1:40, depending on initial cell density. Fresh hPSC medium was added the following day, and the cultures were maintained with routine passaging using this protocol.

### hPSCs preservation

For long-term storage, cell lines were cryopreserved in clumps using Bambanker (Lymphotec Inc / Fujifilm 302-14681) and frozen in CoolCell freezing containers (Corning 432000) at −80 °C. After 24 hours, cryovials were transferred to liquid nitrogen for long-term storage. Thawing was performed by briefly incubating cryovials in a 37 °C water bath until ∼90% of the contents were thawed. The cell suspension was diluted in hPSC medium with 2 µM Thiazovivin, centrifuged at 200 × g for 3 minutes, and the resulting pellet was resuspended in hPSC medium with 2 µM Thiazovivin and plated into three wells of a 6-well plate (80% of the pellet in the first well, 15% in the second, and 5% in the third) to allow for optimal recovery. After 16-18 hours medium was replaced with fresh hPSC medium.

### hPSCs quality controls

Master Cell Banks (MCBs) were established for both lines at passage numbers below 20, and Working Cell Banks (WCBs) were generated below passage 30. Experimental work was conducted using cells that underwent no more than 15 passages after thawing from a WCB vial. MCBs and WCBs were tested for genomic stability using G-banding (resolution: 425 bands, no. of cells karyotyped: 4) and SNP-array (resolution: 1.5Mb). MCBs and WCBs authenticated via STR (15 loci plus amelogenin) and SNP-array match, tested for mycoplasma and tested for sterility. MCBs and WCBs were tested for morphology and expression of markers of undifferentiated state (OCT4, NANOG, SSEA-4 and TRA-1-60) by immunofluorescence and flowcytometry using the following antibodies: anti TRA-1-60 (BD Biosciences, 560173), anti SSEA-4 (BD Biosciences, 560218), anti OCT4 (Santa Cruz Biotechnology, sc-5279) anti NANOG (BD Biosciences, 560791). Differentiation potential for the MCBs was measured using the STEMdiff Trilineage Differentiation Kit (Stem Cell Technologies, 05230) following manufacturer’s recommendations. RNA was isolated from the differentiated and undifferentiated cells at day 5 (endoderm and mesoderm) and day 7 (ectoderm). Markers expression was assessed via RNA sequencing.

### Plasmids

The following plasmids were obtained from Addgene and used in this study. pMDLg/pRRE (Plasmid #12251)(Dull *et al*., 1998), RSV-Rev (Plasmid #12253)(Dull *et al*., 1998), pCMV-VSV-G (Plasmid #8454)(Stewart *et al*., 2003), pFUW-tetO-RUNX1 (Plasmid #139821)(Rosa *et al*., 2018), Teto-BRN3A (Plasmid #62221)(Blanchard *et al*., 2015), Super PiggyBac Transposase (System Biosciences PB210PA-1). TetO-FUW-NGN2-Puro, TetO-FUW-NGN1-Puro, TetO-ISL1, and TRE3G-BI-NGN1-ISL1-Puro were cloned and the maps are available upon request.

### Lentivirus packaging

HEK/293T cells were cultured in MEF media. On the previous day of transfection 10 cm plates were coated with 5 mL of 15 μg/mL polyornithine (Sigma) in Dulbecco’s phosphate buffered saline (DPBS), incubating at 37 °C for 1 h, and washing with PBS twice. 7 × 10⁶ cells of HEK/293T cells, dissociated using 0.25% Trypsin-EDTA (Gibco), were seeded in 6 mL MEF media per plate. After 24 h, the medium was changed to fresh MEF media and cells were transfected using a polyethylenimine (PEI)-based method. DNA (20 µg total: 10 µg of lentivirus plasmid, 5 µg of pMDLg/pRRE, 2.5 µg of RSV-Rev, 2.5 µg of pCMV-VSV-G) was mixed with 500 µL DMEM (Tube A), while 60 µL of 1 mg/mL PEI 25K (Polysciences Inc.) in DPBS was mixed with 500 µL DMEM (Tube B). Tubes A and B were combined and incubated for 20 min at RT, then added dropwise to the cells (1mL per plate). 6 h post-transfection, cells were washed with PBS and prewarmed MEF media was added. Supernatants were collected at 30 and 46 h post-transfection, filtered through a 0.45 µm filter. The virus was concentrated by ultracentrifugation at 21,000 rpm for 2 h at 4 °C. The resulting pellet was resuspended in DMEM/F12 (contains HEPES) in a volume 100 times smaller overnight. Then same amount of 1M sucrose in DMEM/F12 was added and stored at −80 °C after being aliquoted.

### Generation of hPSC lines with PiggyBac-integrated transgenes

WTC11 were seeded at 300–400 × 10³ cells per 3 cm dish in StemFlex medium supplemented with TV and cultured overnight. On day 1, cells were transfected with one PiggyBac plasmid: TRE3G-BI-NGN1-ISL1-Puro and Super PiggyBac Transposase — using Lipofectamine Stem Transfection Reagent (Invitrogen). Transfection complexes were prepared by mixing Lipofectamine and plasmid DNA at a ratio of 1:5 (and maintaining a transposase:vector ratio of 1:2) in warm Opti-MEM I reduced serum medium (Gibco); after a 10 min incubation at RT, 250 µL of the mixture was added dropwise to cells pre-equilibrated in 2 mL warm Opti-MEM I reduced serum medium. Following a 4 hour-incubation at 37°C, 2 mL of warm StemFlex medium was added, and the cells were cultured overnight. On day 2, the medium was aspirated and replaced with 1.5 mL of fresh warm StemFlex medium. Selection was initiated on day 3 by switching to StemFlex medium supplemented with 2 µg/mL puromycin (Millipore Sigma). The selection period was maintained for 4–6 days with medium changes every other day. Cells were then passaged once in StemFlex medium containing both puromycin and TV to ensure uniform exposure to puromycin and eliminate false-positive cells.

### Neural induction using forced expression of TFs with lentivirus on hPSC (one-step)

On day -1 WTC11 cells in 6-well plate with PSC medium were detached by being incubated with Accutase (STEMCELL technologies) for 5 min at 37°C and resuspended with PSC medium containing TV and lentivirus carrying TetO-FUW-NGN2-Puro or TetO-FUW- NGN1-Puro for transfection. Lentivirus was used at a volume of 0.5 µL per well in a 6-well plate. The virus was prepared alongside a TetO-FUW-GFP construct, which consistently yielded >70% GFP-positive cells under the same infection conditions. Based on this, the MOI was estimated to be approximately 10. The cells were seeded in each well of 6-well plates, coated with 1mL of 150 μg/mL Geltrex in DMEM/F12. From day 0, the medium was switched to N3 medium, prepared mixing the followings; 485 mL of DMEM/F12, 5 mL of N2 supplement (Thermo Fisher Scientific), 5 mL of NEAA, 10 mg of Insulin (Sigma), and 5 mL of Pen Strep. And Doxycycline (Dox, Sigma) was added from day 0 to 6 to activate NGN1 or NGN2 expression. Puromycin (Millipore Sigma) was used for selection on day 1 and 2. On day 7, The cells were detached by being incubated with accutase for 20 min at 37°C and resuspended with Neurobasal B27 medium containing TV. Neurobasal B27 medium was prepared by mixing the followings; 483 mL of Neurobasal (Thermo Fisher Scientific), 5mL of Glutamax (Thermo Fisher Scientific), 10 mL of B27 (Thermo Fisher Scientific), 2.5 mL of Pen Strep (Thermo Fisher Scientific). The cells were seeded in each well of 6-well plates, coated with 1mL of 150 μg/mL Geltrex in DMEM/F12. After that the cells were maintained in Neurobasal B27 medium until the day of the analysis.

### Neural induction using forced expression of TFs with lentivirus or PiggyBac on NCCs (2-step)

On day -1 WTC11 cells or and KOLF2.1J cells were detached by being incubated with accutase for 5 min at 37°C and resuspended with PSC medium containing TV. The cells were seeded in each well of 6-well plates, coated with 1mL of 60 μg/mL Geltrex in DMEM/F12. On day 0, the medium is changed to KSR medium; 192.5 mL of Knockout-DMEM-F12 (Thermo Fisher Scientific), 50 mL of KO serum replacement of KSR (Thermo Fisher Scientific), 2.5 mL of Glutamax (Thermo Fisher Scientific), 5mL of NEAA (Thermo Fisher Scientific), and 2 μL of 14.3M 2-Mercaptoethanol (Sigma). The medium is gradually transitioned from KSR to Neurobasal B27 from day 0 to day 11. From day 0 to 7, 100 µM of LDN193189 (Tocris Bioscience), and 10 μM of SB431542 (Tocris Bioscience) and from day 4 to day 11, 10 μM of DAPT (Tocris Bioscience), 3 μM of CHIR99021 (Tocris Bioscience), and 10 μM of SU5402 (Tocris Bioscience) are added to the culture medium.

On day 11, the cells were detached by being incubated with accutase for 20 min at 37°C and resuspended with Neurobasal B27 medium containing TV. The cells were seeded in each well of 6-well plates, coated with 1 mL of 150 μg/mL geltrex in DMEM/F12. Lentivirus carrying TetO-FUW-NGN1-Puro or TetO-FUW-NGN2-Puro, TetO-ISL1, TetO-BRN3A, or pFUW-tetO-RUNX1 are added as described in each experiment. Dox was added from day 12 to 18 induce gene expression. Puro was added on day 13 for selection for lentivirus-transfected cells. 25 ng/mL of human BDNF (R&D Systems) and 25 ng/mL of human GDNF (R&D Systems) were added from day11 to 18. For RNA-extraction, 25 ng/mL of human β-NGF (PeproTech) was added instead of BDNF and GDNF from day 18 to the day of analysis, day 21 (for the experiment for Figure S2C, β-NGF was not added and BDNF and GDNF were kept added in one group). For immunofluorescence staining (IF) and MEA, on day 18, the cells were detached by being incubated with accutase for 20 min at 37°C and resuspended with Neurobasal B27 medium containing TV again. 100K cells were seeded with 20K CD-1 IGS mouse glia cells onto coverslip coated with 100 μL of 150 μg/mL Geltrex in 24 well plate for IF and 24 or CytoView MEA 48 plates coated with 20 μL of 150 μg/mL Geltrex for MEA. From the day after this replate, the cells were cultured in Neurobasal B27 medium with 25 ng/mL of human β-NGF and 2% FBS (Gibco) to the day of analysis (for the experiment for Figure 4B, S2B and S2C, β-NGF was not added and BDNF and GDNF were kept added in one group). From day 28, 4 μM of Cytosine β-D-arabinofuranoside hydrochloride (Ara-C, Sigma) was added.

### RNA extraction and quantitative real-time PCR

Total RNA was extracted using the Phasemaker Tubes (Thermo Fisher Scientific) in combination with TRIzol Reagent (Thermo Fisher Scientific). Briefly, samples were lysed and homogenized in TRIzol Reagent following the manufacturer’s protocol. After chloroform (Sigma) addition, the lysates were centrifuged at 12,000 × g for 5 min at 4°C to achieve phase separation. The aqueous phase containing RNA was carefully transferred.

RNA was precipitated with isopropanol (Sigma), washed with 75% ethanol (Sigma), and resuspended in RNase-free water. For cDNA synthesis, 200 ng to 1 µg of RNA was reverse transcribed using the High-Capacity cDNA Reverse Transcription Kit (Thermo Fisher Scientific), following the manufacturer’s instructions. The resulting cDNA was stored at −20°C until use. Quantitative real-time PCR was conducted using the SYBR master mix (Thermo Fisher Scientific). The reactions were carried out in 384-well plates on an Applied Biosystems QuantStudio 5 instrument. The sequences of primer employed included: *GAPDH*, *MAP2* (PrimerBank (Spandidos *et al*., 2010) ID: 87578393c1), *NTRK1* (PrimerBank ID: 56118209c1), *PRPH* (PrimerBank ID: 66932907c1), *TRPV1* (PrimerBank ID: 117306163c1), *IL31RA* (PrimerBank ID: 336391154c1), *OSMR* (PrimerBank ID: 270288819c1), *HRH1* (PrimerBank ID: 149158708c1), NPPB (PrimerBank ID: 83700236c1), *JAK1* (PrimerBank ID: 102469033c1), *P2RX3* (PrimerBank ID: 332078493c1), *RET* (PrimerBank ID: 126273513c1), *SST* (PrimerBank ID: 71979669c1).

For each experiment, *GAPDH* was utilized as housekeeping control. The rt-qPCR data were analyzed using the comparative CT method. Gene expression levels were normalized to *GAPDH*, and relative expression was determined using the level in PSC (normalized to *GAPDH*) as the reference value (set to 1) for group comparisons.

### Bulk RNA-seq

For RNA library preparation Poly(A) selection was used to enrich for mRNA, followed by sequencing on an Illumina platform with a 2×150 bp configuration, aiming for 50 million paired-end reads per sample. Quality control was performed using FastQC (v0.12.1, https://www.bioinformatics.babraham.ac.uk/projects/fastqc/), ensuring that all samples met the thresholds for basic statistics, per-base sequence quality, and per-sequence quality scores. Adapter sequences and low-quality bases were removed using Trim Galore (v0.6.10, https://www.bioinformatics.babraham.ac.uk/projects/trim_galore. The cleaned reads were aligned to the GRCh38 genome assembly using STAR (Dobin *et al*., 2013) with the GENCODE v29 annotation to generate BAM files and gene count data. Transcript abundance was quantified using Kallisto (Bray *et al*., 2016) with GENCODE v29 annotation and the GRCh38 genome assembly. To calculate TPM values, Kallisto abundance estimates were used and aggregated to the gene level using transcript-to-gene mapping derived from GENCODE annotations. Log-transformed TPM values (log_10_(TPM + 1)) were computed for normalization and subsequently standardized to Z-scores for comparative visualization. For the heatmap of selected genes, the standardized (Z-scores) log-transformed TPM values were used to visualize the expression patterns of predefined genes of interest across samples. This provided a focused view of key gene expression dynamics. For clustering to assess overall similarity between groups, DESeq2 (Love *et al*., 2014) was employed to identify differentially expressed genes (DEGs) via a likelihood ratio test (LRT). Significant DEGs (p-adj < 0.05) were used for hierarchical clustering and heatmap visualization.

Pairwise differential expression analysis between two groups (N1, N2 and N1+I vs N1+B, N1+R, N1+I vs N1+B, or N1+I vs NI + R) was performed using DESeq2, with an adjusted p-value threshold of 0.05. Volcano plots were generated using the DESeq2 output. Ontology enrichment and tissue-specificity analyses of the DEGs were performed with Metascape (https://metascape.org/), using TOP 300 DEGs in each group.

### Multielectrode array (MEA)

Using the Maestro Pro (Axion Biosystems), the activity of the cells was typically measured for 10 min in a 37°C and 5% CO_2_ atmosphere. Recordings were taken at a sample rate of 12.5 kHz/channel with a Butterworth band-pass filter (200 Hz–3000 Hz). Positive and negative deflecting spikes were detected using an adaptive threshold method set at 6x standard deviation of the noise floor. Stimulation assays were performed primarily on day 42 of differentiation; when the same culture batch was reused, wells were rinsed twice with PBS immediately after recording and allowed to recover for at least 3 days in fresh medium before the next stimulation experiment. For stimulation experiments, after baseline measurements, a 10-fold higher concentration of reagent than the final concentration was quickly added to the existing culture solution at 1/9 of the volume and measured for 10 min after addition. Stock solutions were stored in PBS or DMSO at even higher concentrations, with PBS and DMSO being less than 0.1% of the total solution at the time of measurement. The following substances were used for stimulation: 1 mM of Histamine (Sigma), 300 ng/mL of hIL-31 (R&D), 300 ng/mL of hIL-13 (R&D), 10 μM of (E)-capsaicin (Tocris). For the JAK1 inhibition experiment, ABT-317 (provided by AbbVie) was added at a final concentration of 1 μM and incubated at 37°C for 1 h, followed by the addition of hIL-31 at a final concentration of 300 ng/mL. 10 min before and 10 min after the addition of hIL-31 were recorded. The analysis was conducted using Neural Metric Tool (all Axion Biosystems). Mean firing ratio at 10 min post-administration, normalized to 1 for the Mean Firing Ratio at 10 min before administration, was used for comparison between groups. Outliers were identified and excluded using the inter-quartile-range (IQR) method: any well whose mean firing rate fell below Q1 − 1.5 × IQR or exceeded Q3 + 1.5 × IQR was removed prior to statistical analysis.

### Immunofluorescence Staining

Cells were washed with PBS+++ (PBS supplemented with 4 g/100 mL sucrose, 50 µL 0.5M CaCl2, and 50 µL 1M MgCl2), and fixed with 4% paraformaldehyde (PFA) containing 4% sucrose for 5 min at RT. After washing three times with PBS+++, cells were incubated with 0.2% TritonX for 5-10 minutes at room temperature to permeabilize the membranes.

Blocking was performed with blocking buffer (PBS containing 4% BSA, 1% FCS, and NaN3) for 2 h at RT. The cells were then incubated overnight at 4°C with the following primary antibodies diluted in blocking buffer: [anti-Tubb3 (Biolegend, 801201, 1:1000), anti-MAP2 (Abcam, AB5392,1:20000), anti-TRPV1 (Almone Lab, ACC-030,1:200), anti-Trka (Almone Lab, ANT-018, 1:200), anti-IL31RA (Abcam, AB113498, 1:200), anti-OSMR (LSBio, LS-B11477-50, 1:200), anti-RET (Novus biologicals, AF1485, 1:200)].

After incubation, cells were washed three times with PBS and incubated for 1 h at RT in the dark with secondary antibodies diluted in blocking buffer: [Alexa Fluor(AF) 488-conjugated Anti-Chicken IgY (H+L) (Invitrogen, 1:500), AF555-conjugated Anti-Rabbit IgG (H+L) (Invitrogen, 1:500), AF555-conjugated Anti-Mouse IgG(H+L) (Jackson Immunoresearch, 1:1000), AF647-conjugated Anti-Goat IgG(H+L) (Jackson Immunoresearch, 1:500), AF555-conjugated anti-Mouse IgG2a (Biolegend, A-21137,1:500), AF488-conjugated anti-Mouse IgG1 (Biolegend, #A-21121,1:500)]. Cells were then washed three times with PBS, and 0.2ng/mL of DAPI (Thermo Fisher Scientific) was added in the final wash for nuclear staining. Coverslips were mounted using ProLong Gold Antifade Mountant with DNA Stain DAPI (Thermo Fisher Scientific), and imaging was performed using a DMI4000 B Automated Inverted Microscope (Leica Microsystems).

### Mice

CD1 mice (Charles River 022) were housed in specific-pathogen-free (SPF) conditions and environmentally controlled animal facilities with a 12-hour light-dark cycle and were given unrestricted access to food and water. All animal protocols and experiments were approved by the Institutional Animal Care and Use Committee (IACUC) of ISMMS. All experiments were performed following strict IACUC guidelines.

### Mouse glia cell preparation

CD1 pups (postnatal P1-P3) were euthanized via hypothermia-induced anesthesia followed by humane euthanasia on ice, in accordance with institutional animal care guidelines.

Cortices were dissected in ice-cold 10mM HEPES (Gibco) in HBSS (Gibco), and meninges were removed. The tissue was then digested with 80 µL of papain (Worthington), activated with 0.5 μM CaCl₂ (Sigma) and 1 μM EDTA (Invitrogen), at 37°C for 15 min. Following digestion, cells were triturated and filtered through a 70 µm strainer before centrifugation at 1000 rpm for 5 min. The resulting cell pellet was resuspended in mouse embryonic fibroblast (MEF) medium, prepared with 435 mL DMEM (Invitrogen), 50 mL calf serum (Thermo Scientific), 5 mL Pen Strep (Thermo Fisher Scientific), 5 mL sodium pyruvate (Invitrogen), 5 mL Non-Essential Amino Acids (NEAA; Thermo Fisher Scientific), and 4 µL of 2-Mercaptoethanol (14.3M, Sigma). Cells were seeded in a 10 cm dish, with the medium changed after 24 hours. After 5–6 days in culture, cells were detached, resuspended in Bambanker (Lymphotec Inc / Fujifilm 302-14681) and cryopreserved in liquid nitrogen until use.

### Lead contact

Further information and requests for resources and reagents should be directed to and will be provided by the lead contact, Samuele G. Marro.

### Materials availability

Plasmids and cell lines generated in this study will be provided upon request to Samuele G. Marro.

## Acknowledgments

This work was supported by the NIH-NINDS R21NS130319, and Cindy Silvian Foundation Grant to S.G.M. The authors thank the staff of the Stem Cell Engineering Core (SCEC) at the Icahn School of Medicine at Mount Sinai for their technical support and contributions to this study.

## Author contributions

S.G.M., B.K., E.R.G. and P.R. conceived and designed this study; H.I. performed most of the experiments with the help of R.H., X.L. and E.B.; C.H.,I.I., L.M., K.C., L.I.M., and D.Y. contributed data; A.V.H. and K.M.S. analyzed RNA-seq data, S.G.M and H.I. wrote the manuscript with valuable input from all authors.

## Declaration of interests

The authors declare no competing interests.

## Data and code availability

The raw sequencing data for RNA-seq will be deposited in the Gene Expression Omnibus (GEO) repository upon manuscript acceptance and will be available under accession number [to be provided upon acceptance]. Processed data and relevant analysis scripts will be made available upon request. Additional datasets supporting the findings of this study are available from the corresponding author upon reasonable request.

## Abbreviations

hPSC: human pluripotent stem cell
NCC: neural crest cell
TF: transcription factor
AD: atopic dermatitis
CPUO: chronic pruritus of unknown origin
iN: induced neuron
MEA: multielectrode array
DRG: dorsal root ganglion
PB: PiggyBac transposon system

---

**Figure S1.**
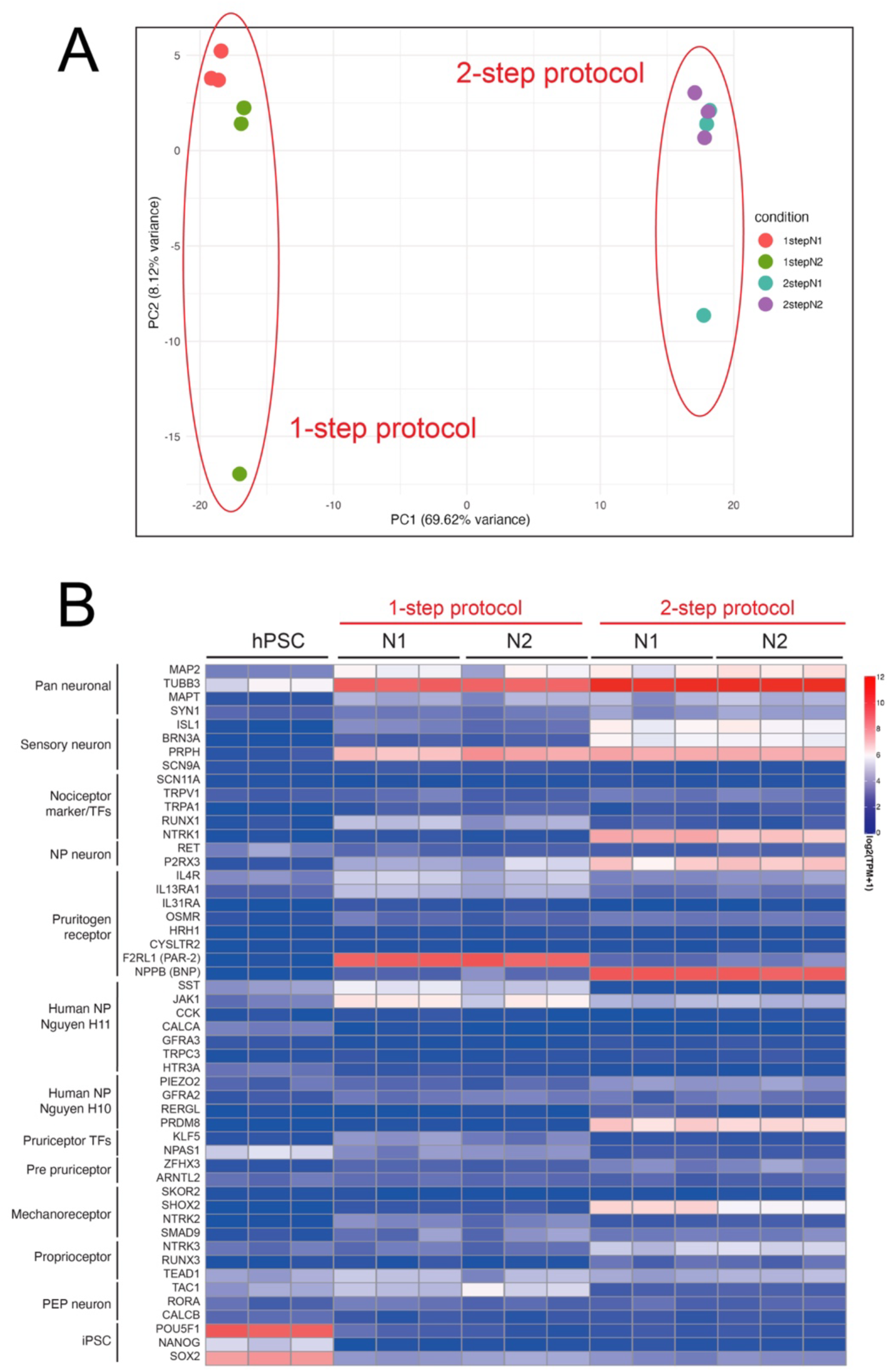
(A) PCA of transcriptomic profiles. Each point represents a sample colored by condition: 1stepN1 (red), 1stepN2 (green), 2stepN1 (cyan), and 2stepN2 (purple). PC1 and PC2 explain 69.62% and 8.12% of the variance, respectively. (B) Heatmap analysis of samples obtained on protocol day 21 using either the 1-step or 2-step protocol with NGN1 or NGN2 expression (1-stepN1, 1-stepN2, 2-stepN1, and 2-stepN2), along with hPSCs, showing the expression of the indicated markers. Transcript expression values (TPM) were log2-transformed, n=3. The analyzed data is same as the one analyzed in Figure 1C.

**Figure S2.**
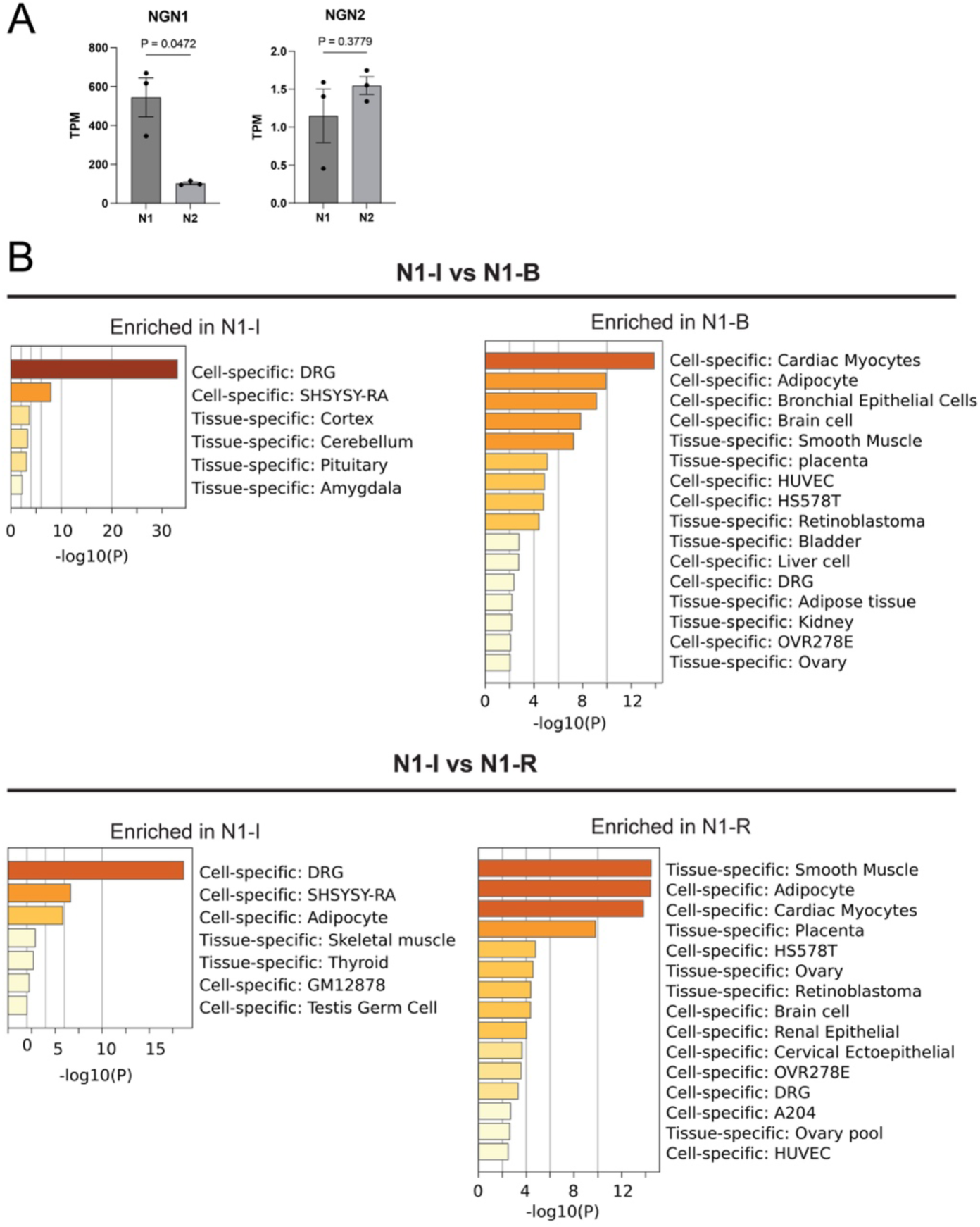
(A) Bar graphs showing TPM of NGN1 and NGN2 in N1 and N2. Each bar represents the mean ± SEM, and individual data points are shown. Statistical comparisons between the two groups were performed using an unpaired t-test with Welch’s correction. (B) Metascape tissue-specific enrichment of the top 300 DEGs: upper panels show 2-step N1-I vs N1-B and lower panels 2-step N1-I vs N1-R; within each comparison, left bars represent tissues/cell types enriched among genes up-regulated in N1-I and right bars those enriched in N1-B or N1-R, with bar length indicating –log_10_ P.

**Figure S4.**
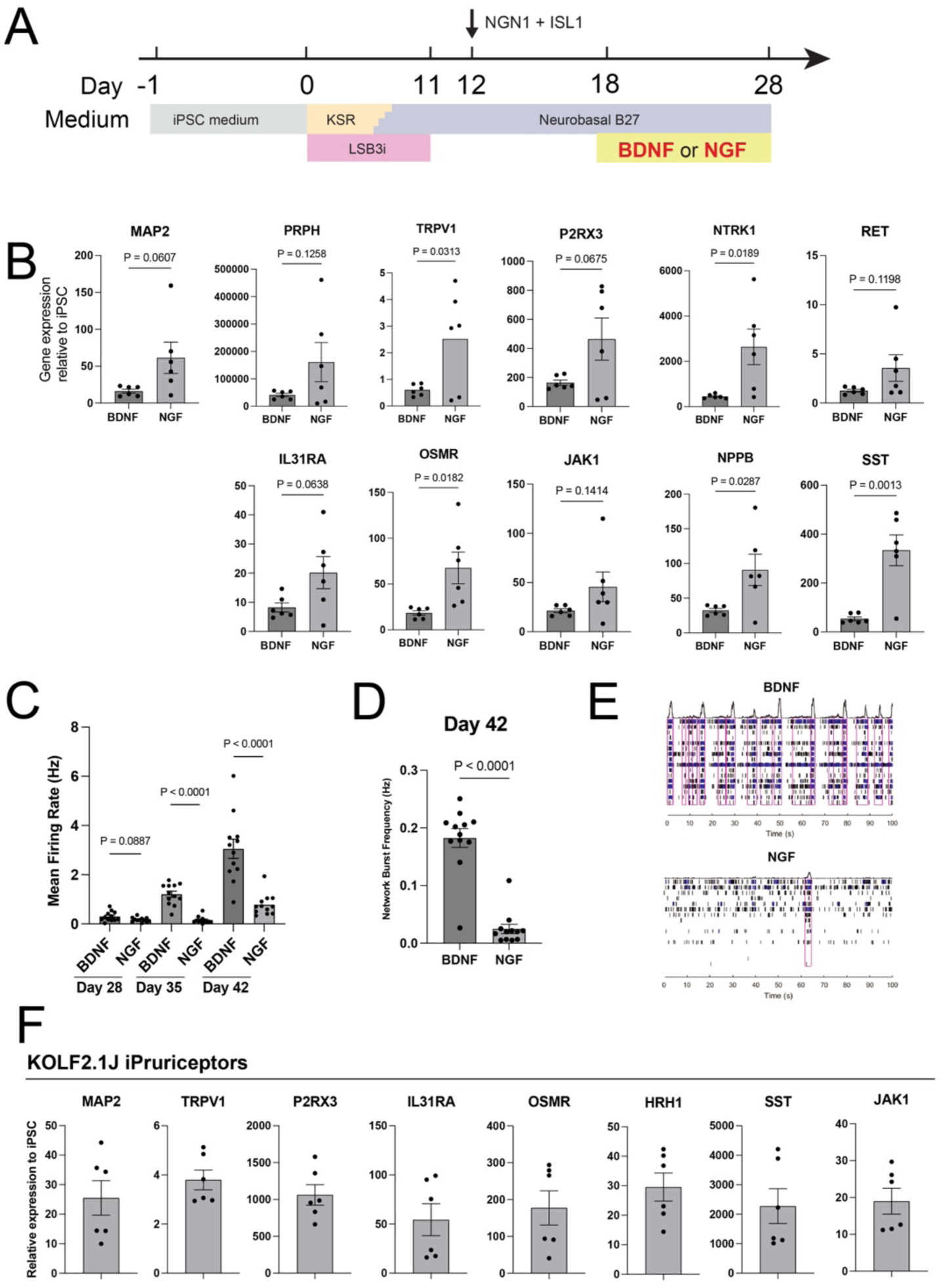
(A) Schematic showing co-expression of NGN1 and ISL1 in NCCs induced by LSB3i via lentiviral transduction, followed by culture with BDNF or NGF from day 18 to day 28, the time point of quantitative PCR analysis. (B) Quantitative PCR analysis of indicated markers on day 28 in the protocols under different neurotrophic conditions (BDNF vs NGF) after day 18. Relative expression levels were normalized to control hPSC levels. Data are presented as mean ± SEM, with individual data points representing independent biological replicates. Statistical comparisons were conducted using an unpaired t-test. (C, D) MEA analysis showing maturation of neuronal activity over time under different neurotrophic conditions (BDNF vs NGF) after day 18. Mean firing rates were measured on days 28, 35, and 42, and network burst frequency was measured on day 42. The values represent the mean firing rate on (C) and network burst frequency on (D). Data are presented as mean ± SEM. Each dot represents one well. Representative data from three independent experiments (n = 12). Statistical comparisons between the two groups were performed using an unpaired t-test with Welch’s correction. (E) MEA raster plots showing neuronal spiking activity of neuronal activity with displaying network burst enclosed in purple box on day 42 under different neurotrophic conditions (BDNF vs NGF) after day 18. Each vertical line represents an action potential recorded from a single electrode. (F) Quantitative PCR analysis of indicated markers on day 21 in the protocols using KOLF2.1J cells. Relative expression levels were normalized to control hPSC levels. Data are presented as mean ± SEM.

## Notes

### Competing Interest Statement

The authors have declared no competing interest.

